# Stroke Shifts Brain Dynamics Toward Criticality: Evidence from Intrinsic Neural Timescales

**DOI:** 10.1101/2025.04.10.647205

**Authors:** Kaichao Wu, Beth Jelfs, Qiang Fang, Leonardo L. Gollo

## Abstract

Stroke disrupts brain function beyond focal lesions, altering multiscale temporal dynamics essential for information processing. We investigated intrinsic neural timescales (INT) and other properties of long-range temporal correlations, using longitudinal fMRI data from 15 ischemic stroke patients across six months, and compared them to age-matched controls. Results show that stroke patients exhibited significantly prolonged INT in multiple cortical regions, reflecting slowed temporal dynamics and disrupted hierarchy. These dynamic changes persisted through recovery and were more pronounced in patients with poor outcomes, especially within cognitive control networks. Computational modelling suggested that stroke-induced INT prolongation driven by heightened neuronal excitability reflects a dynamic shift towards criticality. Our findings position long-range temporal correlations and INT as potential biomarkers for monitoring and predicting functional recovery. This framework provides a novel perspective on stroke-induced brain changes and suggests avenues for targeted neurorehabilitation using interventions aiming at restoring intrinsic temporal dynamics.

## Introduction

The human brain operates through complex dynamics that enable flexible cognition, efficient information processing, and adaptive responses to the environment^1–5^. These dynamics emerge from interactions between neural populations and are crucial for maintaining a balance between stability and flexibility in brain function^6^. As a fundamental property of the dynamics of brain regions, intrinsic neural timescales (INT) reflect the temporal window over which neural activity is integrated^2,3,7,8^. Alterations in INT have been observed across various conditions, including temporal lobe epilepsy^9,10^, Alzheimer’s disease^11^, and Parkinson’s disease^12^, highlighting its potential as a biomarker for disrupted neural dynamics and functional impairments. INT provides valuable insights into how neural architecture shapes functional behaviour and information processing^8,13^, and represents a critical dynamical feature linking brain structure and function. Altered INT are often related^14^ or caused by alterations in gray matter volume^15^, reflecting structural and functional changes in the brain, thereby serving as a bridge between anatomical alterations and dynamic neural processes.

Despite these recent advances in our understanding of INT in different conditions, its role in stroke remains unexplored. As stroke typically causes substantial structural damage to the brain, INT can be a relevant tool to reveal critical changes in brain dynamics. Stroke can result in localized lesions and disruptions to white matter integrity, leading to widespread functional impairments, including deficits in motor control, cognitive processes, and sensory integration^16–18^. While significant progress has been made in understanding post-stroke atypical temporal dynamics in neural activity through studies of network reorganization^19–21^, functional connectivity^22–24^ and network efficiency^25^, the specific impact of stroke lesions on INT and their role in brain function and recovery remains unclear.

Furthermore, INT is derived from the autocorrelation function (ACF) of a neural signal and is measured by its decay properties^3,15,26,27^. A slow decay of ACF, following a power-law function, occurs at a critical point and is known in the literature as critical slowing down^6,28–31^. In practical terms, this means that INT is maximized at a critical point^15^ and various computational properties are also optimized around the critical state^6,28^. This concept of criticality provides a powerful framework for linking INT to distinct dynamical states^15^: subcritical, critical, and supercritical. In the subcritical and supercritical states, INT is short due to rapid (exponential) signal decay; at criticality, they persist longer, following a slow power-law decay. Throughout this framework, criticality has been applied to various brain states^32^, including anesthesia^33,34^, epilepsy^35,36^, neurodegeneration^37^, cognition^38,39^, psychiatry^40^, and sleep medicine^41^. However, its relevance to stroke remains unexplored, presenting an opportunity to investigate how stroke lesions disrupt neural criticality and influence functional recovery. Additionally, since long-range temporal correlations (TCs) have been shown to reflect critical brain dynamics^42–44^, we further examined post-stroke brain dynamics by analysing key TCs properties, including the time delay to reach a specific ACF strength^43,44^ and the Hurst exponent^45^, as complementary measures to INT. We hypothesize that analyzing TCs and INT through the lens of criticality will provide deeper insights into the temporal reorganization of post-stroke brain dynamics and their influence on recovery trajectories.

This longitudinal study examines how stroke disrupts the brain’s multiscale temporal organization by investigating alterations in INT as well as other influential properties of long-range temporal correlations^42–45^ and their hierarchical arrangement across functional networks. Grounded in the criticality framework, we combine longitudinal fMRI with biologically inspired, parsimonious neuronal network modelling to characterize post-stroke temporal reorganization. Empirically, we track INT trajectories during recovery; computationally, we simulate how changes in neuronal excitability reshape brain network dynamics. We test a central hypothesis that stroke generates unique INT signatures that distinguish patients from healthy controls across temporal hierarchies and inform functional recovery outcomes. By bridging empirical findings with computational models of critical dynamics, this work establishes a novel understanding of post-stroke recovery that spans spatial scales, from neuronal excitability to network reconfiguration, while identifying INT as a mechanistic marker of temporal disruption and a potential clinical biomarker for prognosis.

## Results

### Demographic and clinical characteristics

There were no significant differences in age, sex category, and mean framewise displacement between stroke patients and healthy controls (Table I). Patients were first scanned on an average of 23.06 days (standard deviation: 4.32 days) after stroke onset. Recruited patients underwent five follow-up scans at evenly spaced intervals of 30-40 days over six months post-stroke. Recovery trajectories of patients were quantified using the Brunnstrom stage score^46^. We use 2-stage improvements at the final assessment as a cutoff to distinguish the good (n = 6) and poor (n = 9) recovery subgroups^47,48^. There are no significant differences in age or sex categories between the two groups, while the lesion volume is higher in the poor outcome group (3.93±2.24 cm^3^) than in the good outcome group (19.43±19.54 cm^3^). The details of recovery outcome assessment at each follow-up for both groups can be seen in Supplementary Material Table 2.

**Table I:**
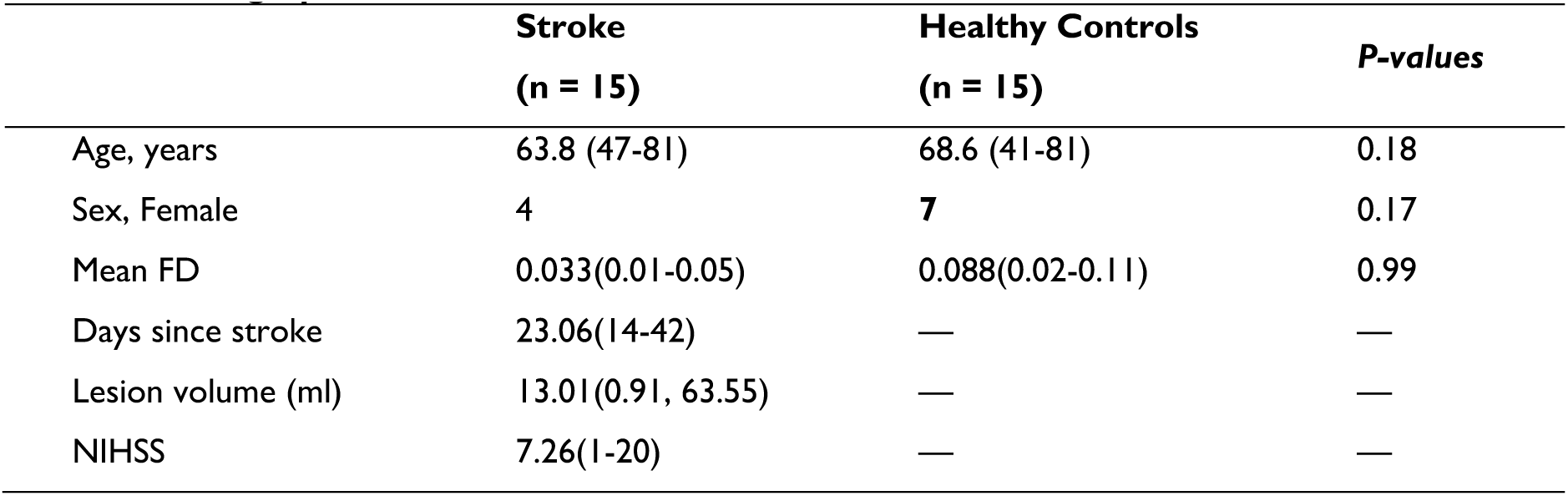
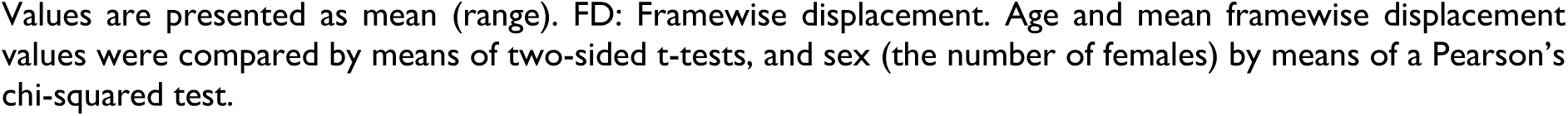
Demographics and Clinical Information.

| | Stroke<br>($n = 15$) | Healthy Controls<br>($n = 15$) | *P-values* |
| --- | --- | --- | --- |
| Age, years | 63.8 (47–81) | 68.6 (41–81) | 0.18 |
| Sex, Female | 4 | 7 | 0.17 |
| Mean FD | 0.033(0.01–0.05) | 0.088(0.02–0.11) | 0.99 |
| Days since stroke | 23.06(14–42) | — | — |
| Lesion volume (ml) | 13.01(0.91, 63.55) | — | — |
| NIHSS | 7.26(1–20) | — | — |

Values are presented as mean (range). FD: Framewise displacement. Age and mean framewise displacement values were compared by means of two-sided t-tests, and sex (the number of females) by means of a Pearson's chi-squared test.

### Abnormal INT Following Stroke

One month after the stroke, stroke patients showed significantly longer intrinsic neural timescales compared to healthy controls, both globally (Figure. 1**D**) and within each specific functional network (Figure. 1**E**). The global average INT was notably higher in stroke patients (*t* = 8.23, *p* < 0.0001, Welch’s t-test, FDR corrected), highlighting widespread alterations in the temporal dynamics of brain activity. When examining specific functional networks, stroke patients showed significant increases in INT across all networks, as seen in the statistics and p-values in Table II.

**Figure 1.**
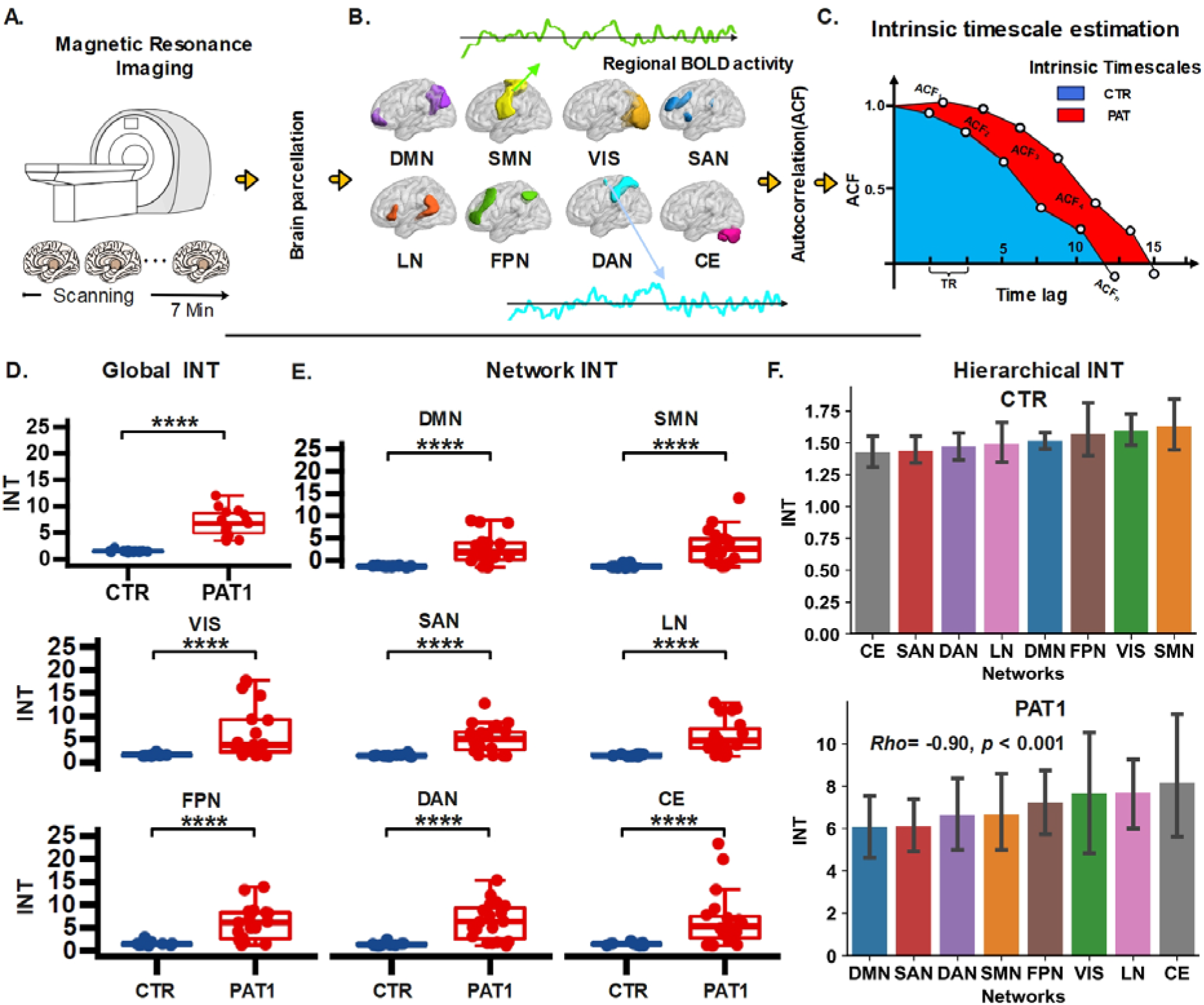
One-month post-stroke analysis of intrinsic neural timescales. **A**. Resting-state fMRI data were collected from a cohort of 15 stroke patients and age-matched healthy controls. Recruited stroke patients (n = 15) underwent five follow-up fMRI scans approximately every 30–40 days over six months post-stroke. **B**. Brain parcellation comprising 32 regions of interest (ROIs) organized into 8 functional networks—Default Mode Network (DMN), Sensorimotor Network (SMN), Visual Network (VIS), Salience Network (SAN), Language Network (LN), Frontoparietal Network (FPN), Dorsal Attention Network (DAN), and Cerebellar Network (CE)— was utilized to extract regional BOLD time series. **C**. Intrinsic neural timescales were estimated by calculating the area under the curve in which the autocorrelation function (ACF) is positive. Compared to healthy controls, one-month post-stroke patients (PAT1) exhibited significantly longer INT both globally, for the whole brain (**D**, *t* = 8.23, *p* < 0.0001, Welch’s t-test, FDR corrected) and within specific functional networks (**E,** see table 1 for detailed statistics). Each dot represents one subject. **F**. The hierarchical Intrinsic Neural Timescales are also disrupted after a stroke, compared to healthy controls (Spearman’s *rho* = −0.90, *p* < 0.0001). The upper and lower error bars display the largest and smallest values within 1.5 times IR above the 75th percentile and below the 25th percentile, respectively. *** indicates p < 0.001 (FDR-corrected); * indicates p < 0.05 FDR-corrected; ns indicates no significance. Abbreviation: INT: intrinsic neural timescales. PAT1: 15 patients recruited and scanned for the first time (first follow-up, approximately 20–30 days post-stroke). CTR: the 15 healthy controls. Data presented in D–F are derived specifically from the first follow-up (PAT1).

**Table 2.** Comparison of INT across all networks between healthy controls and stroke patients.

| Network | T-statistics (CTR < PAT) | p-values |
| --- | --- | --- |
| DMN | 5.92 | < 0.0001 |
| SMN | 5.27 | < 0.0001 |
| VIS | 3.93 | < 0.0001 |
| DAN | 6.82 | < 0.0001 |
| FPN | 6.60 | < 0.0001 |
| LN | 6.72 | < 0.0001 |
| CE | 4.28 | < 0.0001 |

DMN: Default Mode Network; SMN: Sensorimotor Network; VIS: Visual Network; SAN: Salience Network; LN: Language Network; FPN: Frontoparietal Network; DAN: Dorsal Attention Network; and CE: Cerebellar Network. P-values are FDR corrected.

### Reconfigured Hierarchical INT in Post-stroke Patients

In addition to the global and network-specific abnormalities in INT, stroke patients exhibited a significant disruption in the hierarchical organization of INT. Compared with healthy controls (Figure. 1**F**), we found the gradients of mean INT of functional networks are significantly disrupted following stroke. In particular, the CE functional network exhibited the shortest INT (*Mean* ± *Std*: 1.422 ± 0.257) in the control group and the longest INT at PAT1, which corresponds to one month after stroke (*Mean* ± *Std*: 8.178±5.901). Furthermore, the DMN, which had long INT for controls, exhibited the shortest INT in stroke patients (*Mean* ± *Std*: 6.068 ± 2.875). Despite this relative decrease within the stroke group, the INT of the DMN remained longer than that of the controls. Considering all functional networks, the gradient of INT that represents a hierarchical organization of brain dynamics was severely disrupted in stroke patients at 1-month post-stroke (Spearman’s *rho* = −0.90, *p* < 0.0001), as shown in the Figure. 1**F**.

### The Development of INT within Six Months After Stroke

The global average INT was compared between healthy controls and stroke patients across different time points post-stroke to examine the development within a six-month period post-stroke (Figure 2A). Repeated-measures one-way ANOVA analysis revealed significant differences in INT (*F_5,70_* = 13.05, *p* < 0.0001, FDR corrected). The post-hoc t-test indicates that the prolonged INT observed in stroke patients persists for at least five months after the stroke attack (*t_CTR-PAT1_* = 8.22, *p*<0.0001; *t_CTR-PAT2_* = 10.80, *p* < 0.0001; *t_CTR-PAT3_* = 5.94, *p* < 0.0001; *t*_CTR-PAT4_ = 7.61, *p* < 0.0001; *t_CTR-PAT5_* = 5.80, *p* < 0.0001.). This enduring elevation in INT suggests a long-lasting disruption in the brain’s temporal dynamics following a stroke, as depicted in the global INT comparisons. A whole-brain map of the prolonged INT in stroke patients at different time points further highlights the widespread nature of these changes (the example of PAT1 can be seen in Fig. 2B. Other measures of long-range temporal correlations, *INT*_0.1_, *INT*_0.5_ *ACF*_0_, *ACF*_0.1_ and *ACF*_0.5_, further support analogous effects of stroke lesions on the brain’s temporal dynamics (*F*_Hurst_ (*5,70)* = 26.11, *p* < 0.0001, *F*_INT01_(*5,70)* = 9.73, *p* < 0.0001, *F*_INT05_(*5,70)* = 2.69, *p* < 0.0001, *F*_ACF0_(*5,70)* = 26.98, *p* < 0.0001, *F*_ACF01_(*5,70)* = 15.82, *p* < 0.0001, *F*_ACF05_(*5,70)* = 3.38, *p* < 0.01, FRD corrected. See Figure 3).

**Figure 2.**
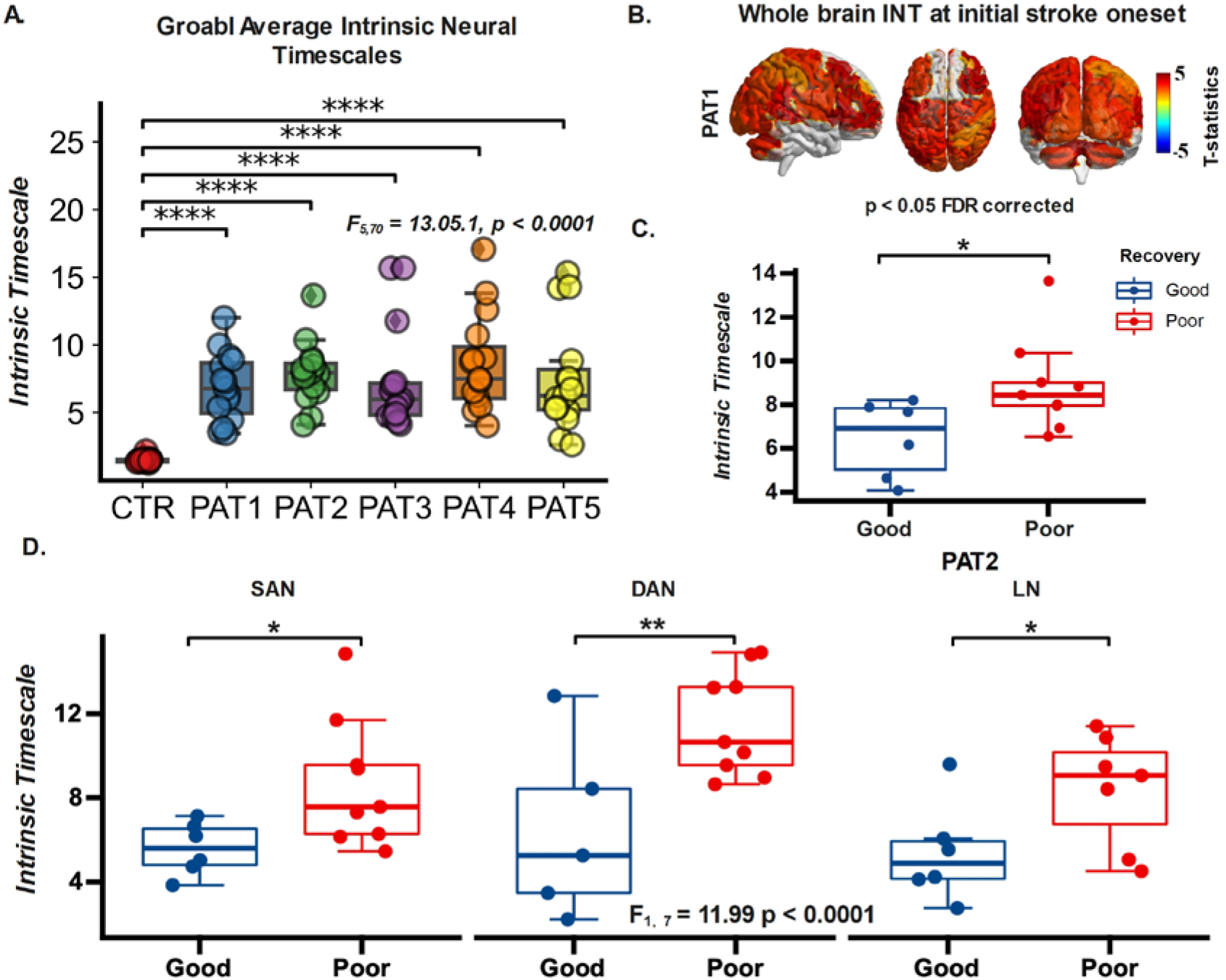
Impact of stroke on intrinsic neural timescales. **A**. Prolonged INT observed in stroke patients persists up to five months post-stroke (*t_CTR-PAT1_* = 8.22, *p*<0.0001; *t_CTR-PAT2_* = 10.80, *p* < 0.0001; *t_CTR-PAT3_* = 5.94, *p* < 0.0001; *t*_CTR-PAT4_ = 7.61, *p* < 0.0001; *t_CTR-PAT5_* = 5.80, *p* < 0.0001, FDR corrected). **B**. Whole-brain map showing prolonged INT in stroke patients at 1-month post-stroke (PAT1). **C**. At 2 months post-stroke (PAT2), significant differences in INT emerged between patients with good and poor recovery outcomes (*t_poor-good_* = 2.30, *p* = 0.03, Welch’s t-test). **D**. Network-level differences in INT between good and poor outcome groups at PAT2. The upper and lower error bars display the largest and smallest values within 1.5 times IR above the 75th percentile and below the 25th percentile, respectively. **** indicates p < 0.0001 (FDR-corrected); ** indicates p < 0.05 FDR-corrected; * indicates p < 0.05 FDR-corrected; ns indicates no significance.

**Figure 3.**
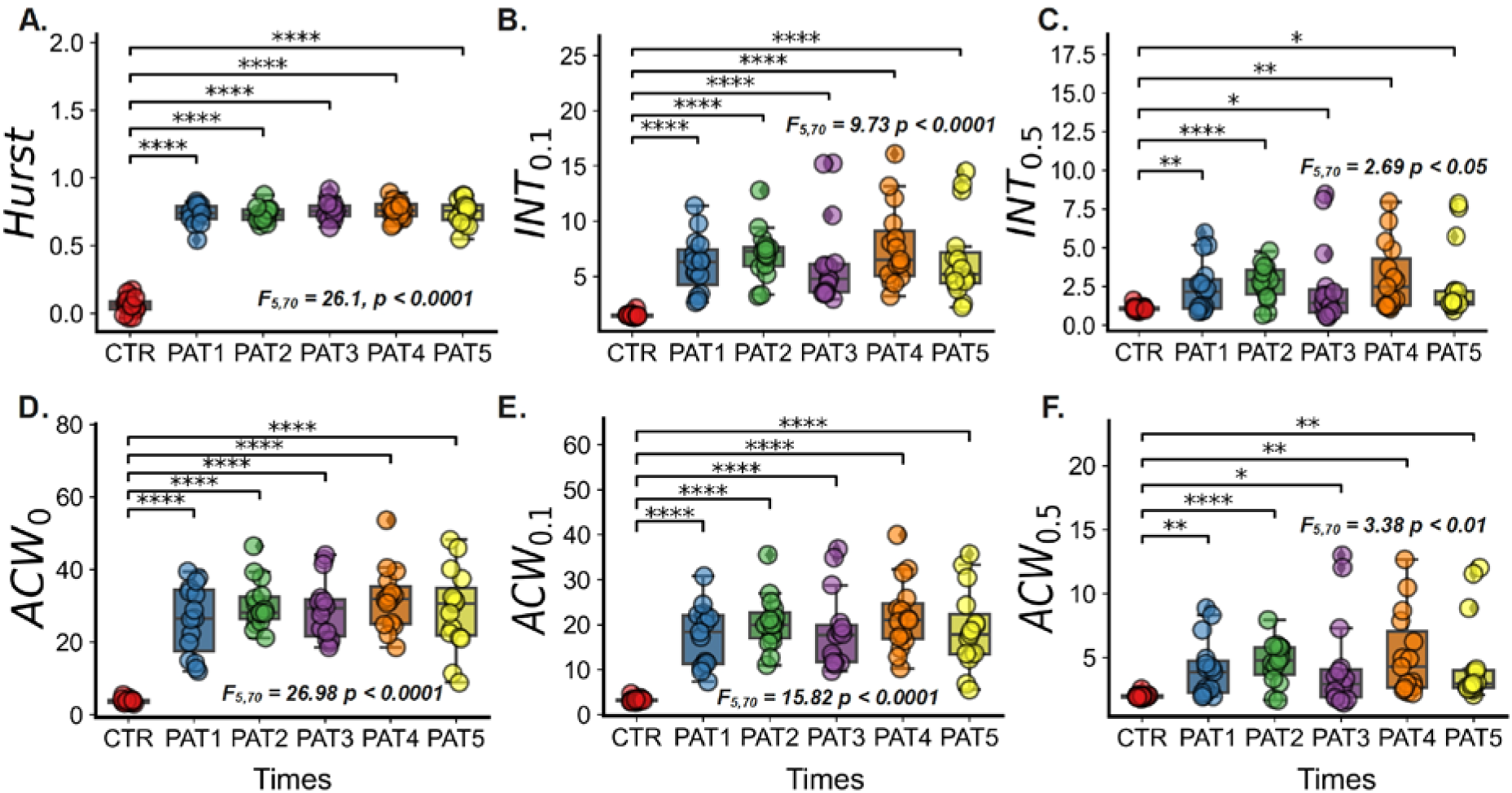
Longitudinal effects of measurements from long-range temporal correlations. A. Hurst exponent, B.*INT*_0.1_, C. *INT*_0.5_, D. *ACW*_0_, E. *ACW*_0.1_ and F. *ACW*_0.5_ sustain the stroke lesions’ disruption on the brain’s temporal dynamics. *F*_Hurst_ (*5,70)* = 26.11, *p* < 0.0001, *F*_INT01_(*5,70)* = 9.73, *p* < 0.0001, *F*_INT05_(*5,70)* = 2.69, *p* < 0.0001, *F*_ACF0_(*5,70)* = 26.98, *p* < 0.0001, *F*_ACF01_(*5,70)* = 15.82, *p* < 0.0001, *F*_ACF05_(*5,70)* = 3.38, *p* < 0.01, FRD corrected. The upper and lower error bars display the largest and smallest values within 1.5 times IR above the 75th percentile and below the 25th percentile, respectively. **** indicates p < 0.0001 (FDR-corrected); *** indicates p < 0.001 (FDR-corrected);** indicates p < 0.05 FDR-corrected; * indicates p < 0.05 FDR-corrected.

### INT as A Prognostic Biomarker for Recovery

To further explore recovery dynamics, patients were categorized into two groups according to motor function outcomes nearly six months later: the good and the poor recovery group. At 2 months post-stroke (PAT2), a significant difference in INT emerged between the two groups (Fig. 2C), with the poor recovery group exhibiting significantly longer INT than the good recovery group (PAT2: *t_poor-good_* = 2.30, *p* = 0.03, FDR corrected), while at other timepoints no significant differences were observed. This finding suggests that differences in INT at 2 months post-stroke may serve as an early predictor of recovery outcomes six months later, highlighting the potential utility of INT as a prognostic biomarker.

A two-way ANOVA was then used at PAT2 to test whether the recovery outcomes (good vs. poor) and the functional networks (DMN, SMN, DAN, etc.) have an effect on INT. We found a statistically significant difference in INT by both outcomes (*F_recovery_* = 11.99, *p* < 0.001, FDR corrected) and by various function networks (*F_networks_* = 3.83, *p* < 0.001, FDR corrected), though the interaction between these terms was not significant. Post-hoc t-test analysis revealed network-specific significant differences in INT between the pairwise recovery groups (Figure. 2D). Patients with good recovery outcomes exhibited notably shorter INT compared to those with poor recovery outcomes in the salience network (SAN; *t_good-poor_* = −3.10, *p* = 0.016), dorsal attention network (DAN; *t_good-poor_* =-5.96, *p* = 0.006) and language network (LN; *t_good-poor_* = −4.31, *p* = 0.031). These findings indicate that lower INT within specific functional networks is associated with better recovery trajectories, further emphasizing the role of INT as a network-level biomarker for predicting post-stroke recovery outcomes.

### Distance to Criticality Explains the Abnormal INT in Stroke

Finally, we interpret the abnormal INT in stroke within the criticality framework. By fixing the network mean degree (*K*) and varying the propagation probability (λ), we control the branching ratio (σ), a key metric that governs neural activity spread and serves as an indicator of criticality. The activity is defined as the instantaneous density of active neurons as a function of time. The examples of neuronal network activities simulated with various σ = [0.08, 0.1, 0.12] with fixed *K* = 10, which represent the subcritical, critical, and supercritical, can be seen in Supplementary Material Figure 2A. The corresponding autocorrelation curves, which are used to compute the INT, are shown in Supplementary Material Figure 2B. As shown in Figure 4A, INT and overall network activity (see Supplementary Material Figure 2C) are directly shaped by σ. Notably, intrinsic timescales increase sharply as the network approaches the critical point (σ = 1), a hallmark of critical slowing down (see Supplementary Material Figure 2D for the single trial relationship between the INT and criticality).

**Figure 4.**
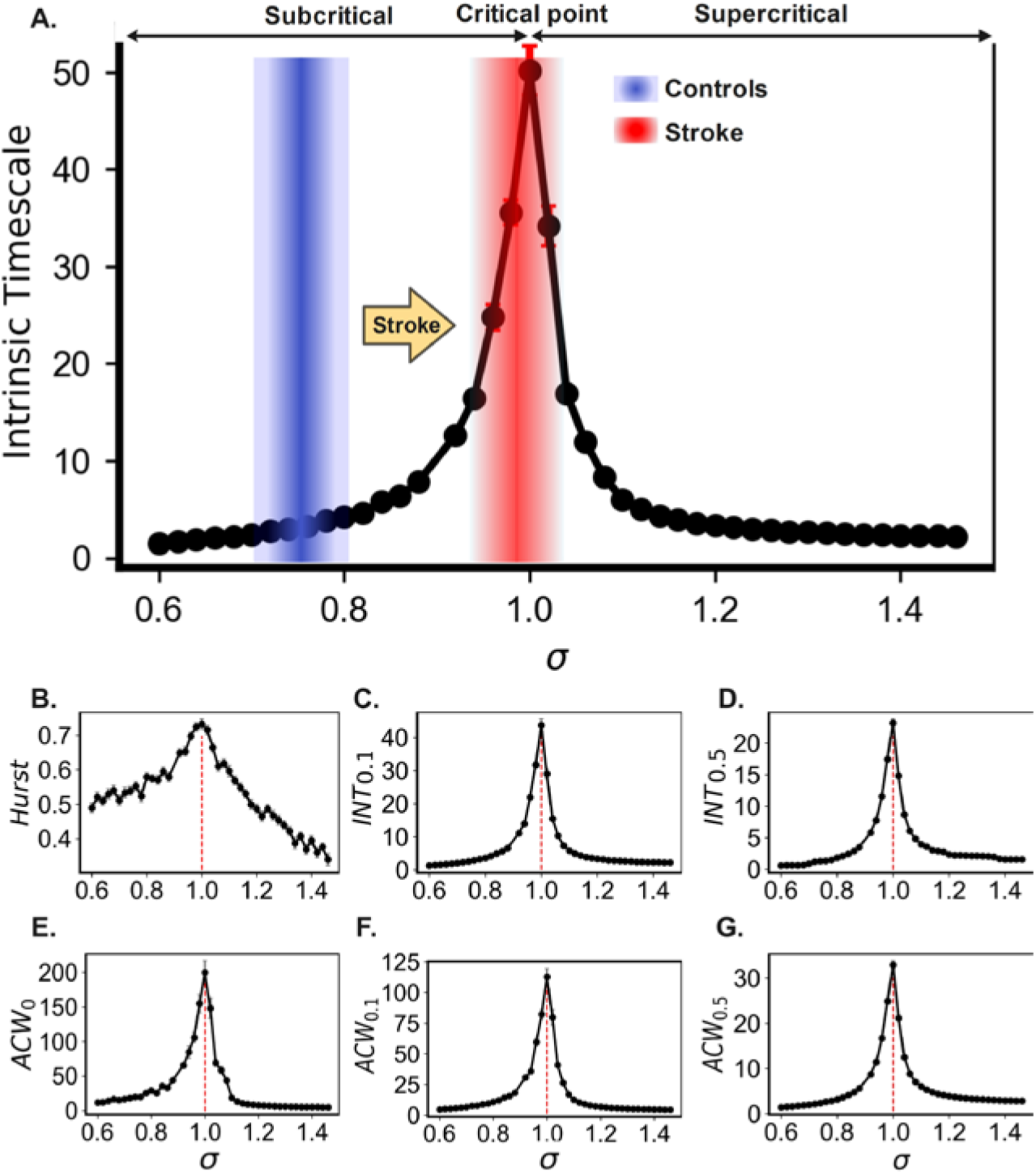
Computation modelling explains how stroke lesions prolong intrinsic neural timescales and alter network dynamics. **A.** The intrinsic neural timescale is shaped by the branching ratio (σ), indicating that stroke shifts the brain dynamics from a slightly subcritical state (blue) toward criticality (red), with the potential to enter a supercritical state. Near a phase transition, cortical network dynamics can be modelled as a branching process, where intrinsic neural timescales peak at the critical point^15^. Stroke-induced shifts toward criticality may eventually drive the system into a supercritical phase, marked by runaway activity that could manifest as seizures. **B–G.** Similarly, complementary measures (see Methods for details)—including the Hurst exponent, *INT*_0.1_, *INT*_0.5_ *ACF*_0_, *ACF*_0.1_ and *ACF*_0.5_—consistently reflect signs of critical slowing down, further demonstrating the robustness and convergence of our findings across multiple temporal metrics. This result is an average of 50 trials. Each dot represents the intrinsic neural timescale (INT) or temporal correlations (TCs) value obtained from the computational model at a given branching ratio (σ). These values were generated by varying the synaptic propagation probability (λ) while keeping network connectivity constant. All networks have *N* = 100,000 neurons and a mean degree *K* = 10 with varied values of λ to satisfy the relationship σ = *K* ⋅ λ. The external driving is given by *r* = 10^−5^. See methods for details.

To further establish the correspondence between our computational model and fMRI signals, we extended the neural simulations and transformed the resulting activity into BOLD-like time series using the canonical hemodynamic response function^49^ (HRF; TR = 2 s). Specifically, we simulated 300s (5 min; time step = 0.001 s) of neuronal activity at different σ values and obtained 150 BOLD-equivalent samples via HRF convolution and downsampling (1 TR). We then computed intrinsic neural timescales (INT; Supplementary Figure 3A–B) and the Hurst exponent (Supplementary Figure 3C–D) from both the raw neuronal dynamics and the HRF-transformed BOLD signals. Although the HRF convolution introduced temporal smoothing and added noise, a peak in both INT and Hurst exponent remained clearly visible near σ = 1 (the critical point). This indicates that the criticality-related modulation of timescales survives the nonlinear transformation into the BOLD domain. While this transformation is not perfect, the inclusion of these HRF-based results (Supplementary Figure 3) provides further support that our model captures relevant temporal features and helps bridge the gap between fast (milliseconds) neuronal-level dynamics and slow (seconds) empirical fMRI observations. Hence, when linked to the prolonged INT observed in stroke patients, these results suggest that stroke increases neural excitability (reflected by λ), shifting brain dynamics closer to criticality and reducing the distance to criticality. Complementary measures related to long-range temporal correlations—including the Hurst exponent, *INT*_0.1_, *INT*_0.5_, *ACF*_0_, *ACF*_0.1_ and *ACF*_0.5_ —further support this interpretation, consistently indicating critical slowing down across multiple estimates (Fig. 4 B-G). This shift renders post-stroke neural networks more susceptible to prolonged activity fluctuations, potentially pushing the system into a supercritical state. Such a transition may drive maladaptive neuroplasticity, contributing to dysfunctional network reorganization and impairing recovery trajectories in stroke patients.

## Discussion

This study investigates the impact of stroke on intrinsic neural timescales (INT) and long-range temporal correlations (TCs) as well as their dynamics during recovery, by analysing functional neuroimaging data from ischemic stroke patients over a six-month post-stroke period. Our findings reveal that stroke induces persistent disruptions in INT and other long-range TCs, both globally and within specific functional networks, reflecting altered temporal dynamics and hierarchical organization. Notably, differences in INT between patients with good and poor recovery outcomes emerge at two months post-stroke, highlighting the potential of INT as a predictor of long-term recovery. The observation of significantly shorter INT in patients with good outcomes further suggests that post-stroke recovery may involve the normalization of INT. These insights advance our understanding of post-stroke brain dynamics and underscore the potential of INT as a biomarker for recovery trajectories.

Changes in INT have been observed across the lifespan, with longer INT in young adults compared to elderly individuals^15^. Atypical INT patterns have also been widely documented in various brain disorders, including shorter INT in sensory regions in autism^14^, mixed patterns in schizophrenia (shorter in parietal and occipital regions^50^ but longer in self-referential processes^51^, and shorter INT in Alzheimer’s disease^11^ and temporal lobe epilepsy^9,10^. Additionally, significant differences in INT have been reported in Parkinson’s disease, with shorter INT in late-stage and longer INT in early-stage patients^12^. Building on this foundation, this study investigates how stroke affects INT and its dynamics during recovery. Our results reveal that stroke patients exhibit significantly elevated INT and long-range TCs compared to healthy controls, with this increase persisting throughout the five-month follow-up period. The observed whole-brain prolongation of INT in stroke patients relative to controls was highly robust and consistent across multiple complementary measures of temporal correlations and the Hurst exponent. All of these measures converged on the same pattern of substantial INT increases in stroke patients. These converging results, together with supporting neurophysiological^52,53^ and modelling evidence^54–56^ of post-stroke hyperexcitability^57,58,59,60^, indicate that the observed INT prolongation is unlikely to be an artifact of the analysis approach and instead reflects genuine alterations in brain dynamics after stroke. The persistent increase in INT suggests enduring disruptions in the temporal integration of neural activity^7,8^, likely reflecting widespread dysfunction in brain network dynamics^19–21^. The consistent elevation of INT across key functional networks—including the default mode network (DMN), salience network (SAN), and frontoparietal network (FPN)—indicates a global impact of stroke. This disruption may impair functional specialization^61,62^ and network efficiency^25^, contributing to the cognitive and motor deficits commonly observed in stroke patients^61,63^.

In addition to elevated INT, our study revealed a significant disruption in the hierarchical organization of INT in stroke patients. The hierarchy INT is consistently observed across both small^13,64,65^ and large-scale fMRI datasets^4,5^. Unimodal regions, such as sensory and motor networks, typically exhibit shorter INT, whereas transmodal regions, including higher-order networks like the central-executive network (CEN), dorsal attention network (DAN), and default-mode network (DMN), tend to display longer INT^5,7,64,66,67^. Our finding of INT in healthy individuals supports and extends the hierarchical organization of INT, demonstrating that temporal dynamics across networks are balanced and optimized for efficient information processing and integration^7,13^. It is worth noting that our network definition was derived from a group ICA map^68,69^, resulting in a relatively coarse parcellation in which large-scale networks such as SMN and VIS may encompass both lower- and higher-order subregions. This spatial averaging across heterogeneous regions can influence the apparent INT hierarchy, potentially explaining why our control group exhibited longer INTs in SMN and VIS compared to the canonical hierarchy reported in previous literature^3,7,70^. Additional methodological factors, such as reliance on resting-state fMRI alone, specific preprocessing choices, and age-matching of the control cohort to the stroke group, may also contribute to these differences. Importantly, in healthy controls, INT hierarchy showed the expected systematic ordering across most networks^5,7,15^, and this organization was strongly altered in stroke patients as indicated by a marked negative correlation in the hierarchy relative to controls (Spearman’s rho = −0.90, p < 0.0001). The cerebellum network, whose function is associated with motor coordination and learning, including balance and posture, as well as other cognitive functions^71^, was affected the most. This likely reflects the importance of the cerebellum network in the reconfiguration of brain dynamics involving motor and other cognitive functions. In a similar vein, during this reconfiguration, the default mode network (DMN), which typically exhibits long INT^13^ and stable dynamics^72,73^, also demonstrated significant changes in its dynamics, showing the shortest INT and the least stability among functional networks in stroke patients^74^.

A key finding of this study is the divergence in INT trajectories based on recovery outcomes^16,48,63,74^. At two months post-stroke, patients with poor recovery exhibited significantly longer INT than those with good recovery. This suggests that early-stage INT measurements could serve as a predictive marker for long-term outcomes. In addition, this finding also led to the hypothesis that good recovery could be associated with a normalization of the altered INT, i.e., a network reduction in the magnitude of INT (for example, salient network and dorsal attention network), whereas poor recovery could be associated with an inability to promptly recover the abnormal INT to normal levels. These findings align with the potential of the temporal profile of the responses to transcranial magnetic stimulation observed in EEG signals following stroke^75^. Nevertheless, noted that even among these patients, INT values did not return to the levels observed in healthy controls. This suggests that full recovery of intrinsic temporal dynamics may unfold over a much longer timescale than six months. Stroke-induced disruptions to excitability and network topology may take extended periods to stabilize and reorganize, especially within higher-order transmodal networks. Therefore, longer-term longitudinal studies are essential to fully characterize the trajectory of INT recovery and validate its prognostic utility in clinical practice.

Our findings provide new insights into the mechanisms underlying altered intrinsic neural timescales (INT) after stroke, integrating empirical evidence with computational modelling. Post-stroke increases in INT observed in our data may reflect both pathological hyperexcitability^57,58,59,60^ and adaptive network reorganization^19,21^. From a neurophysiological perspective, stroke can induce pronounced changes in synaptic plasticity—such as long-term potentiation (LTP) and homeostatic adjustments—which modulate network excitability and alter the propagation of neural activity^57^. These plastic mechanisms influence the emergence of network hubs and stabilize global activity patterns, both of which are linked to proximity to criticality. In our modelling framework, the synaptic propagation probability (λ) serves as a proxy for these excitability changes. This idea is associated with previous modelling studies, which have demonstrated that resting-state networks emerge naturally near criticality when large-scale dynamics are constrained by empirical structural connectivity^76^. Also, empirical research has demonstrated that the intrinsic neural timescales, derived from the BOLD signal, reflect neural activity integrated over long timescales corresponding to the slow fluctuations (<0.1 Hz) captured by resting-state fMRI^15,76^. Our results extend this principle to the stroke context, suggesting that post-stroke increases in excitability can push the system toward a more critical state, thereby lengthening INT and potentially enhancing integration capacity in reorganized networks. This interpretation fits well with infra-slow scale-free dynamics recently discussed by Ao et al.^77^, in which criticality and scale-free dynamics form a background dynamic state that modulates foreground temporal processing, such as INT. From this perspective, stroke-induced changes in excitability may disrupt the background scale-free regime, thereby altering INT.

Increasing λ effectively mimics enhanced recurrent excitation, consistent with recent studies showing that longer INT in the cortex is associated with stronger local recurrent connectivity and elevated baseline activity^78,79^. An increase in λ represents an increase in excitability which functionally parallels the effect of enhanced local recurrent excitation, thereby sustaining activity and prolonging integration windows. In this sense, stroke-related increase in INT may reflect a shift toward stronger effective recurrent excitation in reorganized networks, consistent with the hyperexcitability documented in post-stroke physiology^57,58,59,60^. By varying λ while holding network structure constant, we could systematically shift the model between subcritical, critical, and supercritical regimes, observing that INT peaked at criticality and declined with distance to criticality (DTC). This behaviour aligns with theoretical predictions and prior modelling work on critical brain dynamics^15,43^. Notably, such excitability changes may arise not only from acute neurochemical alterations but also from progressive intra-regional structural remodelling following stroke^80,81^, including changes in synaptic reorganization^82^, local recurrent connectivity^83^, and circuit rewiring^84^. These processes, which unfold across several weeks to months, may sustain or even exacerbate effective excitability within local networks, contributing to prolonged INT. This interpretation is consistent with recent findings linking INT to recurrent synaptic structure^78,79^, and highlights the value of INT as a dynamic marker of both physiological state and ongoing plasticity in the context of stroke recovery^56,82,85^. Nevertheless, structural remodelling can also alter the effective structural degree *K*, which— together with λ—determines the branching ratio (σ = *K*·λ) and thus the DTC. Such *K*-driven shifts can move dynamics toward or away from σ ≈ 1 and thereby modulate INT. In the current computational framework, we varied λ while holding *K* fixed; future work relaxing this constraint and allowing *K* to vary should capture the network reorganization more explicitly.

The relationship between stroke and intrinsic neural timescales can be understood through the framework of criticality^6,28,32,86,87^, which posits that the brain operates near a transition point between ordered and disordered states to benefit from its optimal computational properties. Shifts in the distance to criticality have been applied to various clinical contexts, including depression^30,88^, schizophrenia^89,90^, epilepsy^91,92^, insomnia^92^, Alzheimer’s disease^93^, Parkinson’s disease^94^, and also states like aging^15,95–98^, anesthesia^99^, meditation^100^, cognitive tasks^101^, sleep homeostasis^102^, and sustained wakefulness^103^. Intrinsic neural timescales, and other properties of long-range temporal correlations, serve as a proxy for the brain’s proximity to criticality. Stroke disrupts this balance through mechanisms such as excitotoxicity, inflammation, and maladaptive plasticity, often resulting in hyperexcitability and an increased risk of seizures^104–106^. Post-stroke brains exhibit prolonged long-range temporal correlations compared to healthy controls, indicating a shift toward criticality (see Figure. 4) and potentially into a supercritical state, where pathological dynamics with excessive synchronization can emerge. This shift aligns with the criticality model, in which long-range temporal correlations, commonly used to measure the distance to criticality^107^—peak at criticality and decay in both subcritical and supercritical regimes^15^, linking post-stroke changes in temporal dynamics to network instability. To further contextualize these findings within the framework of brain criticality, the complementary metrics of long-range temporal correlations, including the Hurst exponent and autocorrelation-based indices, were examined^42–44^. These analyses consistently revealed elevated temporal correlations in stroke patients, reinforcing the interpretation that stroke shifts brain dynamics toward a more temporally persistent and potentially critical regime.

Notably, INT is unevenly affected across the brain, causing alterations in the hierarchy of timescales, as some brain regions are more affected than others. This disruption in the hierarchical organization of timescales further highlights how stroke-induced changes can lead to an imbalance in the brain’s functional networks. Thus, the criticality framework provides a unifying perspective on stroke-induced disruptions, translating the complexity of molecular and vascular processes into their net effect on neural excitability. However, variability in stroke location and severity complicates the clinical quantification of deviations from criticality in individual patients. Despite these challenges, criticality remains a valuable conceptual tool for understanding how stroke alters brain network dynamics.

Our findings suggest that the prolonged long-range TCs observed in stroke patients may reflect a maladaptive shift in neural dynamics, driven by excessive network persistence and impaired information processing. The observed association between shorter INT and improved recovery highlights the potential of modulating INT as a therapeutic target for functional restoration.^8^ Given the strong link between INT, neuronal excitability, and network interactions, our results indicate non-invasive brain stimulation (NIBS) techniques—such as transcranial magnetic stimulation (TMS) and transcranial electrical stimulation (TES)—as promising strategies to normalize aberrant INT^1,8,108–113^. Specifically, excitatory stimulation (e.g., anodal transcranial direct current stimulation, high-frequency TMS, intermittent theta burst stimulation, and entrainment by transcranial alternating current stimulation^114^) could be used to reduce abnormally prolonged INT, aligning with the dynamics observed in patients with better recovery outcomes^113^. By targeting influential hub regions or larger brain systems with altered INT, NIBS offers a novel therapeutic avenue to restore balanced network dynamics, enhance functional recovery, and improve post-stroke outcomes. Future research integrating real-time neurophysiological monitoring and computational modelling could further refine these interventions^110^, enabling personalized stimulation protocols tailored to individual neurophysiological profiles. This approach holds significant promise for optimizing recovery trajectories and advancing precision medicine in stroke rehabilitation.

This study provides valuable insights into the neural dynamics of stroke recovery and underscores important avenues for future exploration. The longitudinal design, featuring five follow-ups over six months, provides a robust framework for investigating INT in stroke patients and could be extended to refine recovery trajectory models. The sample size (N=15) provides meaningful insights and establishes a strong foundation for future large-scale studies to further validate and expand upon these findings. Besides, the findings of this study are based on stroke patients with heterogeneous lesions; hence, stronger findings are expected in a more homogeneous group. In terms of modelling, while our approach leveraged the Kinouchi– Copelli framework^115^ to simulate shifts in network excitability via the propagation probability λ, it did not explicitly include inhibitory neuron populations. This simplification allows a clear definition of criticality through the branching ratio, but omits the biological specificity of excitation–inhibition (E/I) dynamics. Given that stroke is known to disrupt inhibitory circuits and GABAergic signaling^116^, future models incorporating distinct excitatory and inhibitory units could offer a more mechanistic understanding of how E/I imbalance contributes to altered INT and recovery potential. Additionally, the focus of the study on fMRI data opens the possibility for integrating multimodal approaches, such as EEG or diffusion-weighted imaging, that could deepen our understanding of the temporal and structural changes associated with INT dynamics. The identification of meaningful patterns of INT across functional networks marks a notable advancement. Future research utilizing advanced techniques, such as machine learning or personalized network analyses, holds significant potential to reveal individual-level variability and improve predictive accuracy. While this study focused on motor recovery, future research could explore the impact of INT and TCs on cognitive performance, particularly in functions associated with the salience, language, and dorsal attention networks, which showed significant predictive value for recovery outcomes.

## Materials and Methods

### Subjects

Fifteen ischemic stroke patients (ISP) admitted to the First Affiliated Hospital of Shantou University Medical College were recruited for this longitudinal study. The recruited patients were followed up in six months across five timepoints. Inclusion criteria were as follows: (i) The first stroke or previous stroke without sequelae; (ii) The diagnosis was in line with the main points of cerebrovascular disease diagnosis approved by the fourth National Cerebrovascular Disease Academic Conference in 1995; (iii) The onset time was 2 weeks to 6 months, and Brunnstrom stage 0-3 of the affected hand; (iv) No metal implants or pacemakers in the body; (v) No previous history of epilepsy; (vi) Informed consent signed by the patient or his immediate family member. Participants were excluded if they met any of the following conditions: (i) Patients with Parkinson’s disease or other neurological diseases; (ii) Patients with serious diseases of the heart, lung, and other organs; (iii) A history of mental illness, drug abuse, and alcohol abuse; (iv) Patients with an unstable condition, such as blood pressure index: low-pressure ؾ90 mmHg or high-pressure >160 mmHg.

The fifteen patients presented with mild to severe motor function deficits (National Institutes of Health Stroke Scale, NIHSS: *mean*: 7.26, 1-20). Among these patients, 5 had right-sided, and 10 had left-sided strokes, with an average age of 63.81 years (*standard deviation*: 11.68 years). The corresponding lesion map of all stroke patients has been attached to Supplementary Material Figure 1. The cohort consisted of 4 males and 11 females, with the first MRI scan performed on average 23.06 days post-stroke (*standard deviation*: 4.32 days). Ethical approval for the study was granted by the Medical Research Ethics Committee of the named hospital (approval number: SUMC-2021-78-K), and all participants provided written informed consent in accordance with the Declaration of Helsinki. Additionally, a control group of 15 age-matched healthy individuals with no history of stroke and a normal neurological examination (7 males, 6 females, mean age: 68.61 years, standard deviation: 6.42 years) was included for comparison. No significant differences were observed between the stroke patients and healthy controls in terms of age (*p* = 0.18, two-sample t-tests) or sex distribution (*p* = 0.17, Pearson’s chi-squared test). All aspects of this study were also approved by the Monash University Human Research Ethics Committee. Demographic features of recruited stroke patients and healthy controls are listed in Table 1.

Each patient underwent five resting-state fMRI scans spaced approximately 30–40 days apart over a six-month period post-stroke. For clarity, we refer to the first, second, third, fourth, and fifth follow-up scans across all patients as PAT1, PAT2, PAT3, PAT4, and PAT5, respectively. In practice, all fifteen recruited patients finished the follow-up plan. Hence, there are 75 scans for 15 patients and 15 scans of 15 healthy controls for this unique longitudinal dataset (90 MRI scans in total). Given the observational nature of the study, treatment regimens were not standardized across participants, allowing for natural variability in clinical care. Recovery trajectories of patients were quantified using the Brunnstrom stage score^46^. This validated measure of upper limb motor recovery function is assessed across four dimensions. To test the association between INT and post-stroke recovery, patients demonstrating at least a 2-stage improvement by the final assessment were classified as having good recovery, while those showing improvement by one or fewer stages were classified as poor recovery^47,48^. The detailed demographic characteristics and the follow-up clinical features of stroke patients can be seen in Supplementary Material Table 2. There was no statistically significant difference in time since stroke (*p* = 0.59, permutation test) and size of lesion volumes (*p* = 0.16, permutation test) between subgroups.

### fMRI Acquisition, Pre-Processing, and Denoising

Resting-state fMRI data were acquired for all patients at five follow-up visits using a 3.0 T Discovery MRI scanner with an 8-channel head coil at the SUMC MRI Center. High-resolution T1-weighted anatomical images were acquired using a multi-planar rapid gradient echo sequence with the following parameters: 129 slices, repetition time (TR) = 2250 ms, echo time (TE) = 4.52 ms. Following the anatomical scan, resting-state functional MRI data were collected using a single-shot gradient-echo EPI sequence: TR = 2000 ms, TE = 30 ms, flip angle = 90°, and voxel size = 3.43 × 3.43×5.0 mm^3^ (no gap). A total of 210 volumes were obtained over a 7-minute duration for each MRI scan (see Figure 1A).

The fMRI data underwent preprocessing using a tailored pipeline within the CONN functional connectivity toolbox^117^, integrated with the Statistical Parametric Mapping software (SPM12)^118^. For fMRI from stroke patients, the bias of the hemisphere was eliminated by flipping the right hemispheric lesions to the left along the midsagittal plane (see Supplementary Material Figure 1C for the left lesions of all patients). For each participant, the first 10 dummy volumes were discarded. The subsequent functional images were corrected for slice timing and head motion (no significant difference in framewise displacement detected, *p* = 0.99, two-sample t-tests). Outlier detection was performed using Artifact Detection Tools^117^ to identify anomalous time points for each participant. These outliers were removed as covariates, and the remaining function images were then normalized to the Montreal Neurological Institute (MNI) space after masking lesioned tissue.

Non-smoothed functional images were further processed through the default denoising pipeline^117^, which included removing confounding effects and applying temporal band-pass filtering. Potential confounding effects in blood oxygenation level-dependent (BOLD) signal include noise components from (i) cerebral white matter and cerebrospinal areas, (ii) estimated motion parameters identified during the realignment step of pre-processing, (iii) outlier scans from outlier identification (i.e., scrubbing), and (iv) constant and (v) first-order linear session effects. While recent ECoG-fMRI studies have demonstrated that the global signal contains meaningful neuronal information related to arousal and neuro-vascular coupling^119,120^, global-signal regression (GSR) can introduce artefactual anticorrelations and potentially remove genuine neural fluctuations^121^. Hence, no global signal regression was applied. We regressed out the five confounds with CompCor algorithm^122^ (a component based noise correction method for BOLD timeseries) implemented in CONN^117^, and then the residual time series was band-pass filtered in the 0.008–0.09 Hz range^117^.

### Head motion control

The head motion effect was controlled by calculating the individual mean and maximum framewise displacements (FD)^123^:

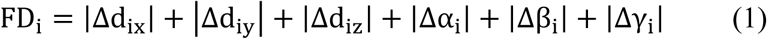

where Δ*d*_*ix*_ = *d*_(*i*−1)*x*_ − *d*_*ix*_ and similarly for other rigid body parameters [*d*_*ix*_, *d*_*iy*_, *d*_*iz*_, α_*i*_, β_*i*_, γ_*i*_]. Participants with a maximum displacement exceeding 1.5 mm and a maximum rotation above 1.5 degrees were excluded. While in practice, no FD was above the threshold, and no subjects were excluded. Besides, 24 motion parameters calculated from the six original motion parameters using Volterra expansion^124^ were regressed out as nuisance covariates.

### Brain Parcellation

To analyse changes in network dynamics resulting from stroke lesions, we utilized a functional brain parcellation derived from CONN’s group independent component analysis (ICA) of the HCP dataset (497 subjects). This parcellation includes 32 regions of interest (ROIs) spanning the entire brain, grouped into eight large-scale functional networks or systems: Default mode network (DMN), Sensorimotor network (SMN), Visual network (VIS), Dorsal attention network (DAN), Salience network (SAN), Frontoparietal network (FPN), Language network (LN), Cerebellar network (CE) (see Supplementary Material Table 2 for the details of eight large-scale functional network and 32 ROI as well as their peak coordinates). ICA spatial maps were applied to each participant’s fMRI BOLD data to extract a representative timeseries for each ROI (see Figure 1B for parcellation map).

### Estimation of Intrinsic Neural Timescales

We estimated intrinsic timescale values for all regional BOLD timeseries. This involves first computing the autocorrelation function (ACF) for each timeseries:

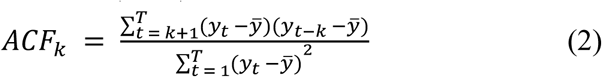

where *y* denotes the preprocessed regional BOLD timeseries, *y^-^* is the mean value across time points. *t* is the length of time bins, which is the time of repetition (*TR* = 2000 ms) of the MRI scan, and *T* is the number of time points (200 in the experiment).

Due to the lower temporal resolution of fMRI data, the intrinsic timescale was calculated as the area under the curve of the autocorrelation function from one to the time lag at which the correlation reaches zero (see Figure 1C for illustration):

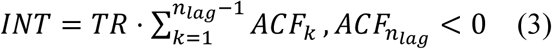

It has been proven that the method proposed to compute the intrinsic neural timescales for fMRI data was highly correlated with those calculated from simultaneously recorded EEG data^14^. This process was applied to all regional BOLD timeseries of patients and healthy controls, resulting in an intrinsic timescale map for the entire brain network for each participant. To further validate potential biases in the estimation of INT across the literature, we incorporated additional metrics commonly used in studies of long-range temporal correlations (TCs). Specifically, we computed alternative INT measures based on the area under the autocorrelation function (ACF) curve up to defined thresholds: the point at which the ACF first decays to 0.5 (denoted as *INT*_0.5_) and 0.1 (*INT*_0.1_)^42,43^. Given the well-established relationship between temporal autocorrelation and the Hurst exponent^125^, we also included the Hurst exponent as a complementary measure of TCs. Furthermore, we quantified ACF strength^43^, defined as the time lag at which the ACF curve first crosses zero after linear interpolation (denoted as *ACW*_0_). To enhance robustness, we additionally computed threshold-based ACF indices—0.5 (*ACW*_0.5_) and 0.1 (*ACW*_0.1_)—corresponding to the time lag above these respective threshold values. These supplementary TCs-based measures serve to cross-validate and reinforce our INT-based findings, offering a more comprehensive characterization of post-stroke temporal brain dynamics. See Table III for the details about Intrinsic Neural Timescales and additional measures of temporal correlations.

**Table 3:**
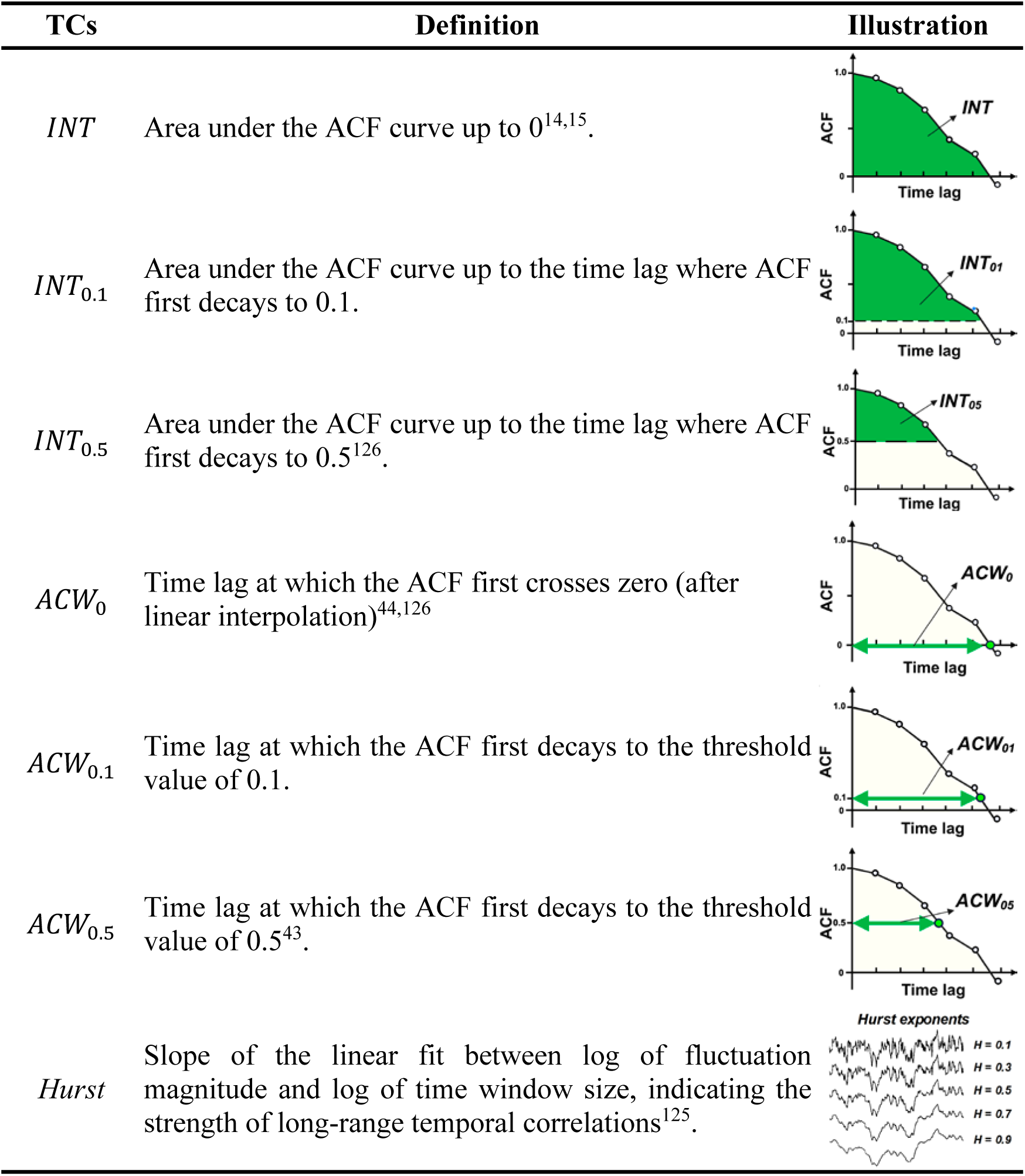
Intrinsic Neural Timescales and Additional Measures of Temporal Correlations.

| TCs | Definition | Illustration |
| --- | --- | --- |
| $INT$ | Area under the ACF curve up to 0<sup>14,15</sup>. | |
| $INT_{0.1}$ | Area under the ACF curve up to the time lag where ACF first decays to 0.1. | |
| $INT_{0.5}$ | Area under the ACF curve up to the time lag where ACF first decays to 0.5<sup>126</sup>. | |
| $ACW_0$ | Time lag at which the ACF first crosses zero (after linear interpolation)<sup>44,126</sup> | |
| $ACW_{0.1}$ | Time lag at which the ACF first decays to the threshold value of 0.1. | |
| $ACW_{0.5}$ | Time lag at which the ACF first decays to the threshold value of 0.5<sup>43</sup>. | |
| $Hurst$ | Slope of the linear fit between log of fluctuation magnitude and log of time window size, indicating the strength of long-range temporal correlations<sup>125</sup>. | |

### Computational Model of Stroke-Induced Network Excitability

To investigate stroke-related alterations in intrinsic neural timescales (INT), we adopted a computational modelling approach grounded in the neurophysiological principles of synaptic plasticity. As reviewed by Stampanoni Bassi et al^57^, long-term potentiation can promote the emergence of highly connected hub regions that support efficient integration, while homeostatic plasticity helps prevent instability by limiting excessive connectivity in peripheral nodes. Stroke disrupts this balance, often inducing local hyperexcitability^104,106^and large-scale network reorganization^19,21^. We hypothesized that such synaptic changes shift the brain’s operating point relative to its critical state, thereby altering its intrinsic timescale structure.

Hence, we implemented a large-scale excitable neuronal network model based on the Kinouchi–Copelli framework^115^ to simulate and explain stroke-related alterations in intrinsic neural timescales (INT). In particular, a random network of *N* = 100,000 excitable spiking neurons and a mean average degree *K* = 10 was utilized to initialize brain neuronal networks. The neuronal network dynamics were then modelled using the Kinouchi-Copelli model^115^. Each neuron is a cyclic cellular automaton that follows a discrete-time process defined on the state space (quiescent, spike, and refractory). When at a quiescent state, neurons can spike by (i) an external drive modelled as a Poisson process with probability with the rate *r* = 10^−5^, which activates neurons with probability ℎ = 1 − *e*^−*r*⋅δ*t*^, where δ_*t*_ = 1 ms is the time step, or by (ii) receiving input propagation from a spiking connected neuron with probability λ. The spiking neuron will enter a refractory state at the next time step and then enter the quiescent state after a period of 8 ms. We set the time length as *T* = 5050, including a 50-ms transient time window, and then the neuronal network activity is defined as the instantaneous density of active neurons as a function of time.

The branching ratio σ (= *K* ⋅ λ) of the neuronal networks represents the average number of spikes generated by each excited neuron in the next time step. The brain is in a critical state when σ = 1, where the network activity propagates in a balanced manner, maximizing information transfer and computational efficiency^6,115,127^. In the subcritical state (σ < 1) activity decays rapidly, leading to diminished neural interactions and reduced information integration. Conversely, in the supercritical state (σ > 1) activity grows excessively. For branching processes, the branching ratio can indicate deviations from criticality, which is known as the distance to criticality (*DTC*), where *DTC* = 1 − σ in subcritical states and *DTC* = σ − 1 in supercritical states. The INT is maximized at the critical state, and it decreases with DTC towards both sub- and supercritical states^15^. Due to the nature of the branching ratio, changes in network degree *K* or the propagation probability λ will affect σ and, thus, cause alterations in INT. The effects of varying *K* and *N* on the relationship between INT and *DTC* were previously explored in the context of aging^15^ and cognitive performance^43^. Given that stroke often results in hyperexcitability^57,58,59,60^ and an increased risk of seizures^128,129^, here we vary λ while fixing *K* for simplicity to examine how stroke-induced changes in neural excitability alter the relationship between *DTC* and INT. In addition, to relate simulated neuronal activity to BOLD signals, we convolved a long stimulated neuronal network activity (300s) with the canonical hemodynamic response function (HRF) as implemented in SPM^118^.

The simulated neuronal activity was generated at a 1 ms time step and then convolved with the HRF, downsampled to a repetition time (TR) of 2 s to yield BOLD-like time series. We computed INT and the TCs on both the raw neuronal activity and the HRF-transformed BOLD signals at each σ to examine whether criticality-related modulations of timescales are preserved in the fMRI domain.

Our decision to vary synaptic propagation probability (λ) while keeping network connectivity (K) constant was specifically intended to capture changes in neural excitability that are characteristic of stroke-induced hyperexcitability. This is consistent with evidence that stroke alters the balance of excitatory and inhibitory transmission and affects both short-term and long-term synaptic plasticity^57^,which in turn modulates the effective propagation of activity across neural networks. These plastic mechanisms influence the emergence of network hubs and the stabilization of network-wide activity, both of which are tightly linked to criticality and timescale dynamics. Hence, such stroke-induced synaptic reorganization can be effectively abstracted by varying λ, which controls the probability of post-synaptic activation in our model. By linking λ to excitability and plasticity, our model bridges microscale synaptic changes with macroscale functional dynamics, reinforcing the relevance of *DTC* and INT as biomarkers sensitive to stroke-induced network alterations.

### Statistical Analysis

Welch’s t-test was performed to investigate if there were statistically significant differences in the whole brain, global average, and network-level INT between healthy controls and patients. Whenever it involves multiple comparisons, the p-values were corrected with the false discovery rate (FDR) method at a corrected α = 0.05. Repeated-measures one-way ANOVA (level of significance *p* < 0.05, FRD corrected^130^) was used to test if INT significantly differed across different time points. Two-way ANOVA was used to test whether the recovery outcome and functional network have an effect on INT. In the case of significant ANOVA results, post hoc t-tests were performed.

## Acknowledgements

This study acknowledges the Shantou University Medical College for the help of patient recruitment and MRI scan acquisition. This work was supported by the Australian Research Council (ARC), Future Fellowship (FT200100942), the Juan De La Cierva (JDC2024-055992-I) and Ramón y Cajal Fellowship (RYC2022-035106-I) from FSE/Agencia Estatal de Investigación (AEI), Spanish Ministry of Science and Innovation, and the María de Maeztu Program for Units of Excellence in R&D, grant CEX2021-001164-M. The funder played no role in study design, data collection, analysis and interpretation of data, or the writing of this manuscript.

## Data availability

The source data for the main figures presented in the paper are attached. The raw MRI scans of patients and controls will be available upon request after signing the anonymous data protocol. The modelling work, including all trials, can be in a code repository (see code availability for details).

## Code availability

The customized Python codes for data and statistics analysis and network modelling are available in the GitHub repository https://github.com/xiaohajiayouo/Stroke-Alters-Intrinsic-Neural-Timescales-and-Their-Hierarchical-Organization.

## Author Contribution

Conceived and designed the work: K.W and LL.G; Data curation: K.W, B.J, and Q.F; Performed the analysis: K.W and LL.G; Writing – review **&** editing: K.W, B.J, and Q.F, and LL.G. All authors reviewed the manuscript.

