## Supplementary material for "Stroke Shifts Brain Dynamics Toward Criticality: Evidence from Intrinsic Neural Timescales": Updated SM (from another version of this preprint: version 2, 2025-09-22)

**Supplementary Table** I – Demographics and clinical characteristics of participants.

| Patient | Sex | Age | Side of Lesion | Lesion location | Lesion Volumes ($CM^3$) | Artery | Day Since Stroke | Timing BS PAT1 | PAT2 | PAT3 | PAT4 | PAT5 | Recovery |
|---|---|---|---|---|---|---|---|---|---|---|---|---|---|
| ***Good recovery group*** | | | | | | | | | | | | | |
| P2 | F | 65 | R | Cerebellum | 0.91 | PICA/SCA | 26 | 2 | 4 | 5 | 6 | 6 | good |
| P3 | M | 54 | L | Thalamus | 3.61 | Thalamic perforators | 42 | 1 | 2 | 3 | 3 | 3 | good |
| P4 | M | 58 | L | Temporal lobe/basal ganglia | 1.53 | MCA | 22 | 4 | 5 | 6 | 6 | 6 | good |
| P11 | F | 78 | R | Posterior limb of internal capsule | 2.39 | MCA | 22 | 1 | 1 | 3 | 3 | 3 | good |
| P13 | M | 47 | L | Precentral/Mid frontal | 6.23 | MCA | 18 | 3 | 4 | 5 | 5 | 5 | good |
| P14 | M | 57 | L | Pallidum | 6.44 | Pallidum | 18 | 1 | 3 | 3 | 4 | 4 | good |
| *Mean±SD* | - | *59.42±9.05* | - | - | *3.93±2.24* | | *-24.66±8.21* | *2±1.06* | *3.16±1.34* | *4.16±1.21* | *4.50±1.25* | *4.50±1.25* | good |
| ***Poor recovery group*** | | | | | | | | | | | | | |
| P1 | M | 59 | R | Temporal/occipit lobe | 63.55 | PCA | 16 | 5 | 5 | 5 | 6 | 6 | poor |
| P5 | M | 63 | R | Thalamus/Hippcocampus | 8.68 | Thalamic perforators | 20 | 3 | 2 | 2 | 3 | 4 | poor |
| P6 | F | 67 | R | Thalamus/cuneus/lingual | 39.81 | Thalamic perforators | 23 | 1 | 1 | 1 | 1 | 1 | poor |
| P7 | F | 48 | L | Thalamus | 0.95 | Thalamic perforators | 21 | 6 | 6 | 6 | 6 | 6 | poor |
| P8 | M | 48 | L | Pallidum | 1.09 | Pallidum | 24 | 6 | 6 | 6 | 6 | 6 | poor |
| P9 | M | 80 | L | Mid Occipital/lingual | 50.51 | PCA | 26 | 1 | 1 | 2 | 2 | 2 | poor |
| P10 | M | 78 | L | Thalamus | 3.12 | Thalamic perforators | 27 | 6 | 6 | 6 | 6 | 6 | poor |
| P12 | M | 81 | L | Hippocampus | 1.04 | PCA | 27 | 1 | 1 | 1 | 1 | 1 | poor |
| P15 | M | 73 | L | Putamen | 5.33 | Pallidum | 14 | 6 | 6 | 6 | 6 | 6 | poor |
| *Mean±SD* | - | *66.33±12.11* | - | - | *19.43±19.54* | - | *22±4.42* | *3.88±2.23* | *3.77±2.29* | *3.88±2.21* | *4.11±2.18* | *4.11±2.28* | poor |

*Abbreviations: R: Right, L: Left, BS: Brunnstrom stage, SD: Standard deviation.PAT1: 30-40 days, PAT2: 60-70 days, PAT3: 90-100 days, PAT4: 120-130 days. PAT5: 150-170 days.*
*MCA: Middle cerebral artery. PCA: Posterior Cerebral Artery. PICA: Posterior Inferior Cerebellar Artery. SCA: Superior Cerebellar Artery.*

**Supplementary Table** II – The parcellation encompasses eight large-scale functional networks, comprising a total of 32 regions of interest (ROIs).

| # | Network | ROI | Coordinates | # | Network | ROI | Coordinates |
|---|---|---|---|---|---|---|---|
| 1 | Default mode network (DMN) | Medial prefrontal cortex (MPFC) | 1,55,-3 | 2 | Visual network (VIS) | Medial | 2,-79,12 |
| | | Precuneus cortex (PCC) | -39,-77,33 | | | Occipital | 0,-93,-4 |
| | | Lateral parietal (LP) (Right) | 47,-67,29 | | | Bilateral (Right) | -37,-79,10 |
| | | Lateral parietal (LP) (Left) | 47,-67,29 | | | Bilateral (Left) | 38,-72,13 |
| 3 | Salience network (SAN) | Anterior cingulate cortex (ACC) | 0,22,35 | 4 | Sensorimotor network (SMN) | Superior | -55,-12,29 |
| | | Anterior insula (AI) (Right) | -44,13,1 | | | Bilateral (Right) | 56,-10,29 |
| | | Anterior insula (AI) (Left) | 47,14,0 | | | Bilateral (Left) | 0,-31,67 |
| | | Borstal prefrontal cortex (RPFC) (Right) | -32,45,27 | 5 | Dorsal attention network (DAN) | bilateral frontal eye field (FEF) | -27,-9,64 |
| | | Rostral prefrontal cortex (RPFC) (Left) | 32,46,27 | | | bilateral frontal eye field (FEF) | 30,-6,64 |
| | | Supramarginal gyrus (SMG) (Right) | -60,-39,31 | | | intraparietal sulcus (IPS) | -39,-43,52 |
| | | Supramarginal gyrus (SMG) (Left) | 62,-35,32 | | | intraparietal sulcus (IPS) | 39,-42,54 |
| 6 | Frontoparietal network (FPN) | Posterior parietal cortex (PPC) (Right) | 52,-52,45 | 7 | Language network(LN) | Inferior frontal gyrus (Right) | 54,28,1 |
| | | Lateral prefrontal cortex (LPFC) (Right) | 41,38,30 | | | Inferior frontal gyrus (IFG) (Left) | -51,26,2 |
| | | Lateral prefrontal cortex (LPFC) | -43,33,28 | | | Posterior superior temporal gyrus (pSTG) (Right) | -57,-47,15 |
| | | Posterior parietal cortex (PPC) (Left) | -46,-58,49 | | | Posterior superior temporal gyrus (pSTG) (Left) | 59,-42,13 |
| 8 | Cerebellar network (CE) | Anterior | 0,-63,-30 | - | - | Posterior | 0,-79,-32 |

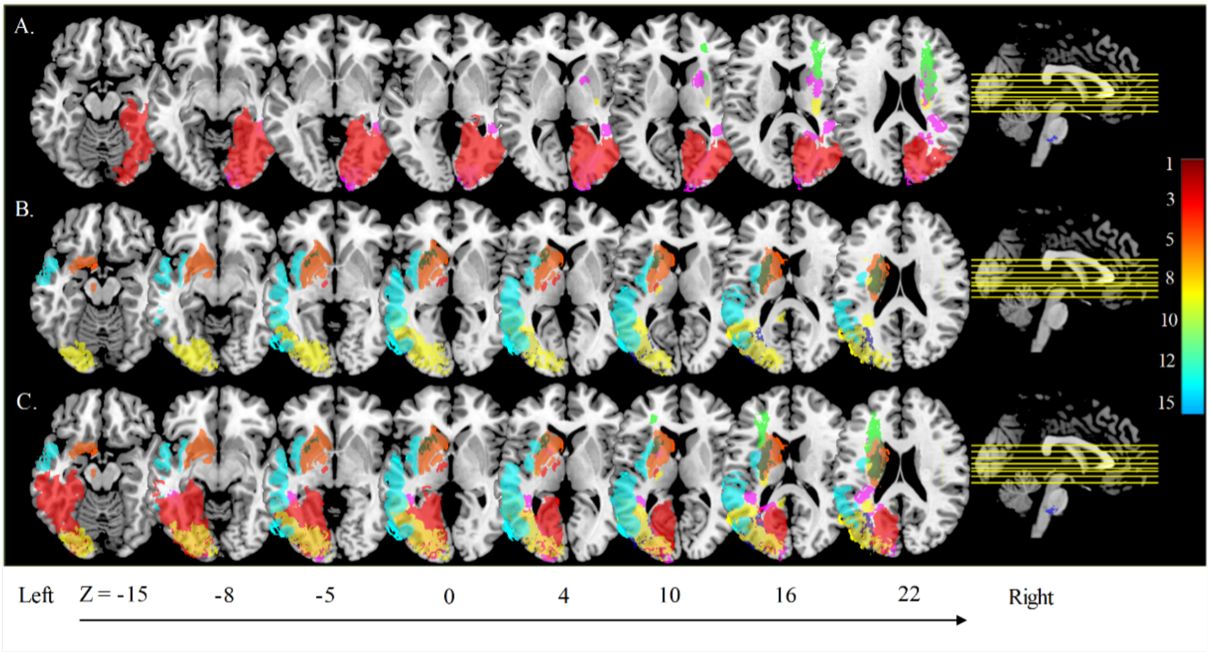


**Supplementary Figure** 1 – **Lesion overlap map across 15 stroke patients**. **(A)** right-sided lesion (n = 5); **(B)** left-sided lesion (n = 10); **(C)** the entire sample of stroke patients with right-sided lesion flipped to left-sided (n = 15). The color bar indicates the patients. See supplementary table 1 for additional details.

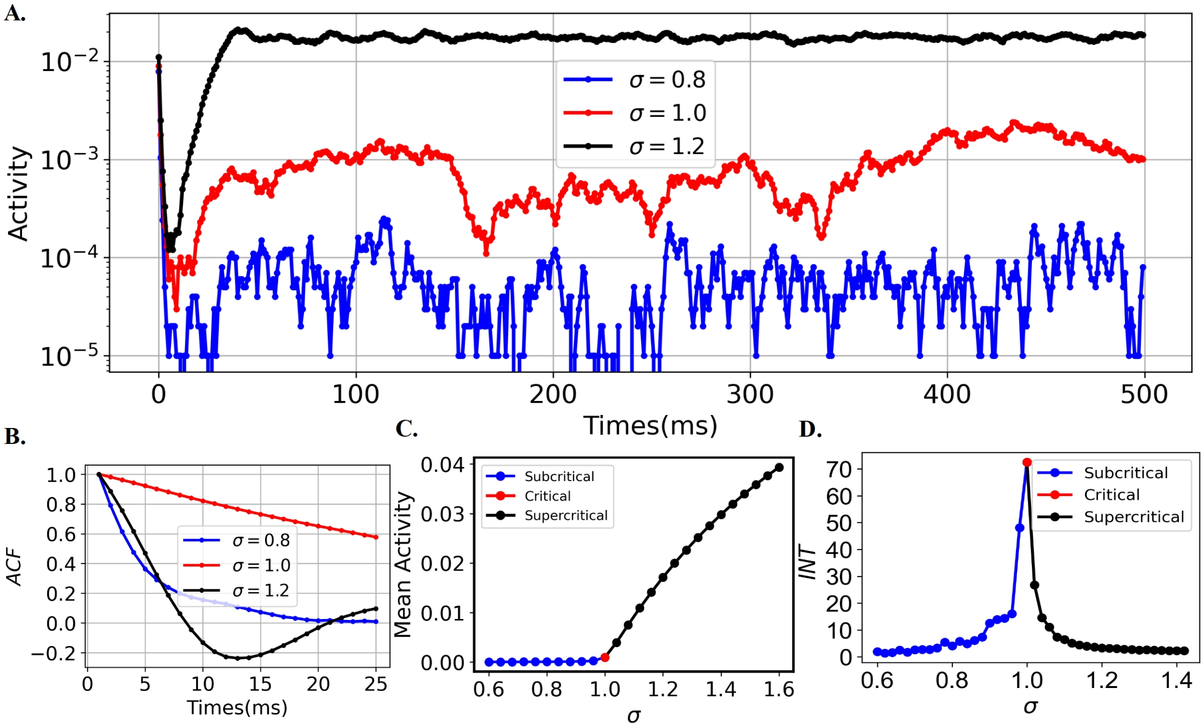


**Supplementary Figure** 2 – **Neuronal network model dynamics.** **(A)** Density of active neurons as a function of time for one trial with branching ratio $\sigma = 0.8$, 1.0, and 1.2. The activity is color coded with blue indicating the subcritical regime, red indicating the critical state, and black indicating the supercritical regime. **(B)** Exemplar autocorrelation function for branching ratios $\lambda = 0.8, 1.0$ and 1.2. **C.** Mean activity as a function of branching ratio. There is a phase transition at $\sigma = 1$. **D.** Relationship between intrinsic timescale and branching ratio for a single trial. All networks have N = 100,000 neurons and a mean degree $K = 10$ with varied values of $\lambda$ to satisfy the relationship $\sigma = K * \lambda$. The external driving is given by $r = 10^{-5}$, see methods for details.

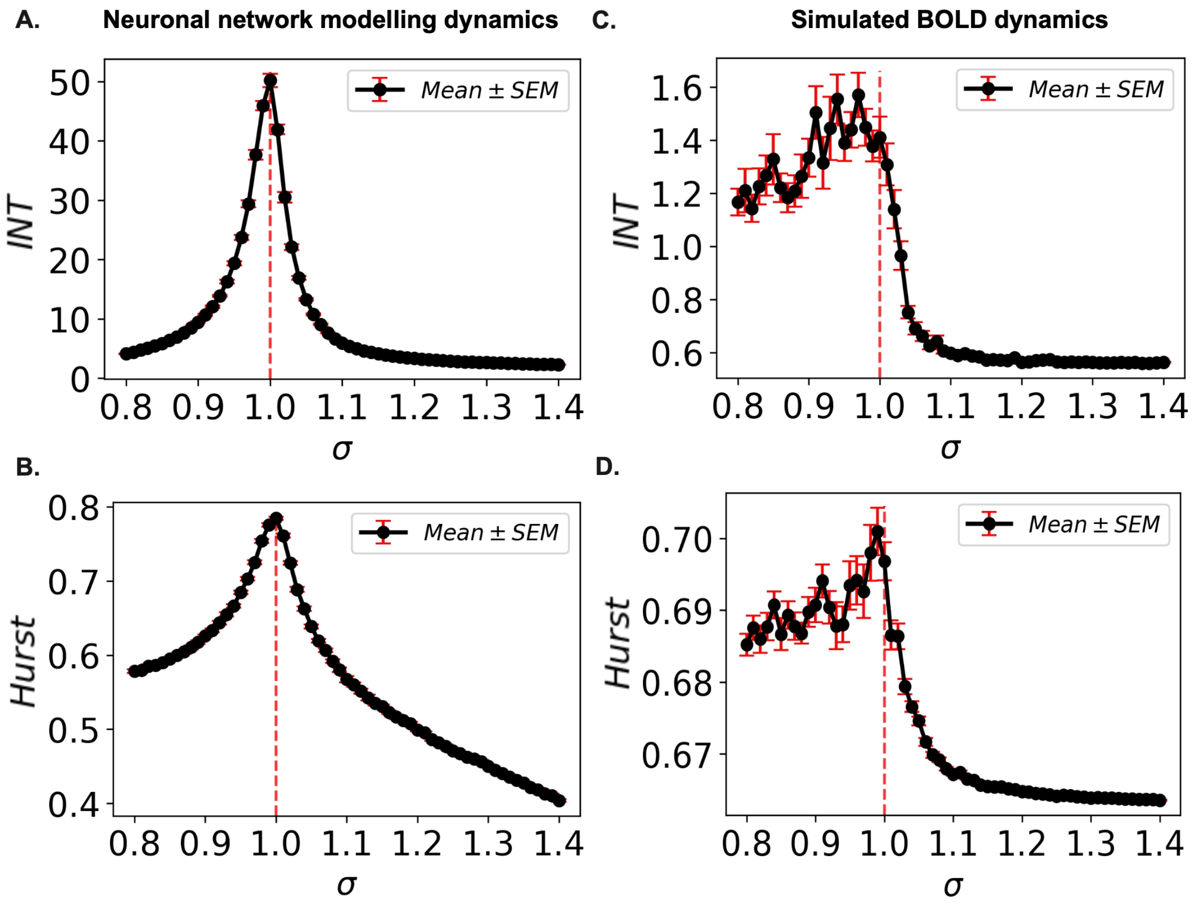


**Supplementary Figure 3 – Signatures of criticality in long simulated neuronal and blood oxygenation level-dependent time series.** Intrinsic neural timescales (INT, **A**, **C**) and Hurst exponents (**B**, **D**) obtained from long simulated time series. The left panels show metrics calculated from 300s of direct neuronal network activity. The right panels show the same metrics from a simulated BOLD signal, generated by convolving the neural activity with a canonical hemodynamic response function (HRF; TR = 2s, 150 timepoints). Results are averaged over 50 trials. Although the HRF convolution attenuates the signal and increases noise, a distinct peak near criticality ($\sigma = 1$) persists in both the neural and BOLD-derived measures.
